# hsa-miR-9-5p highly expressed in syncytiotrophoblast-derived extracellular vesicles from early-onset preeclampsia impairs cerebral microvascular endothelial cell pro-angiogenic capacity

**DOI:** 10.1101/2024.10.21.619546

**Authors:** Prassana Logenthiran, Toluwalase Awoyemi, Gabriel Davis Jones, Maryam Rahbar, Naveed Akbar, Ana Sofia Cerdeira, Carlos Escudero, Manu Vatish

## Abstract

**Background:** Cerebrovascular complications are the leading cause of maternal mortality associated with preeclampsia. Extracellular vesicles (EVs) containing microRNAs (miRNAs) and derived from syncytiotrophoblast (STB-EVs) are suspected to play a role in these complications. Previously, we found that STB-EVs from the placentas of women with preeclampsia have a higher content of the angiogenesis regulator hsa-miR-9-5p. We now investigate the effects of hsa-miR-9-5p on the proangiogenic properties of brain endothelial cells and identify potential protein targets involved in these processes.

**Methods:** Brain endothelial cells (hCMEC/D3) were treated with hsa-miR-9-5p (0, 5 and 10 nM) to assess cell viability and proliferation. Additionally, cell migration and proteomic profile in hCMEC/D3 treated with hsa-miR-9-5p (10 nM) were also analyzed.

**Results:** Compared to control, hsa-miR-9-5p significantly reduced hCMEC/D3 cell proliferation and migration without affecting cell viability. Proteomic analysis identified several vital proteins potentially mediating these effects, including vascular endothelial growth factor type C (VEGFC), placental growth factor (PLGF or PGF), and platelet-derived growth factor B (PDGFB). Treatment with hsa-miR-9-5p did not impair the capacity of hCMEC/D3 to respond to tumour necrosis factor-α (TNF-α).

**Conclusion:** hsa-miR-9-5p reduces hCMEC/D3 cell proliferation and migration, and modulates the expression of angiogenic regulators such as VEGFC, PLGF, and PDGFB, without affecting TNF-α mediated activation of hCMEC/D3. This suggests that STB-EVs cargo hsa-miR-9-5p may selectively inhibit the proangiogenic capacity of brain endothelial cells. These findings enhance our understanding of cerebrovascular alterations in preeclampsia and may guide future studies and therapeutic interventions.

## Introduction

Preeclampsia (PE), characterized by new-onset hypertension and multisystem dysfunction, typically manifests after 20 weeks of gestation. This condition significantly contributes to global maternal mortality rates, mainly attributable to cerebrovascular complications [1-3] with a predominant occurrence observed in low-resource settings [4-6]. Preeclampsia is associated with significant structural and functional alterations in the brain [7, 8]. These changes potentially elevate the risk of both immediate [9, 10] and prolonged adverse outcomes, including a three-fold increase in the susceptibility to vascular dementia, and two-fold increase in the likelihood of stroke [11]. Despite the remarkable relevance for women’s health, the underlying mechanisms of acute and long-term cerebrovascular complications in women who experienced preeclampsia are unknown.

In contrast to organs like the uterus, kidney, and the heart, which undergo heightened perfusion and filtration during pregnancy, the brain must adapt circulation to maintain stability and safeguard its intricate milieu amidst extensive hormonal and cardiovascular shifts [12]. These adaptations encompass structural remodeling of the brain parenchymal microcirculation, involving the development of new brain blood vessels, including arterioles (arteriogenesis), and the formation of new capillaries (angiogenesis) [12]. In the context of preeclampsia, it is conceivable, though currently speculative, that the failure or impairment of new vessel formation in the brain parenchyma may underlie potential acute and long-term cerebrovascular alterations.

While the involvement of the placenta is essential for the manifestation of preeclampsia [13, 14], its specific role in preeclampsia-associated brain complications remains less well elucidated. The placenta under stress in preeclampsia releases soluble factors and syncytiotrophoblast-derived extracellular vesicles (STB-EVs) into the maternal circulation [15-17]. Clinical investigations utilizing plasma samples from women with preeclampsia [18-20] or employing a rat model of preeclampsia induced by placental ischemia [21] have indicated impairments in cerebrovascular endothelial function, suggesting a potential role for circulating placental factors in driving these alterations. Nonetheless, the precise identity of these circulating factors responsible for brain endothelial dysfunction and the mechanisms by which they contribute to impaired new brain blood vessel formation in the context of preeclampsia remain unknown.

Syncytiotrophoblast-derived extracellular vesicles are known to carry diverse cargo, including microRNA (miRNA) [22, 23] and have the ability to reprogram the recipient cells [24]. These STB-EVs can be further delineated into small STB-EVs (< 200 nm, sSTB-EVs) and the relatively less explored medium/large STB-EVs (> 200 nm, m/lSTB-EVs). Notably, the biogenesis and functional roles of these distinct subtypes may carry significant implications. While sSTB-EVs are believed to be constitutively released by cells to facilitate intercellular communication, m/lSTB-EVs appear to be released in response to placental stress [25].

We recently validated differentially expressed miRNAs in m/lSTB-EVs, including high levels of hsa-miR-9-5p (miR-9-5p), isolated from ex vivo dual lobe placental perfusion performed on placentas from women with early-onset preeclampsia (EOPE, < 34 weeks) compared to normal pregnancy (NP) [26]. Furthermore, we observed an upregulation of miR-9-5p within circulating extracellular vesicles (EVs) isolated from serum of these women [26]. miR-9-5p emerged as a candidate of particular interest due to its established role in regulating angiogenesis in pathological conditions such as cancer [27, 28]. Since preeclampsia may involve impairment of brain angiogenesis, we decided to investigate the *in vitro* effect of miR-9-5p on angiogenesis using a human cerebral microvascular endothelial cell line (hCMEC/D3).

## Methods

### Human cerebral microvascular endothelial cell culture

Human cerebral microvascular endothelial cells (hCMEC/D3) utilized in the experiments were provided by Professor Yvonne Couch from the Radcliffe Department of Medicine, University of Oxford. These cells were cultured in complete endothelial cell growth medium-2 (EGM-2) supplemented with the Bullet Kit (Lonza Biologics, Basel, Switzerland) and a 1% v/v penicillin-streptomycin antibiotics cocktail. Cultures were maintained at 37°C in a humidified atmosphere with 20% O_2_ and 5% CO_2_, as previously outlined [20].

### Treatment with miRNAs

hCMEC/D3 were incubated with either EGM-2 growth medium alone, vehicle control (mirVana™ miRNA Mimic, Negative Control #1, ThermoFisher, Waltham, MA, USA), or two concentrations (5 nM and 10 nM) of miR-9-5p (ThermoFisher).

### Cell proliferation and viability assay

hCMEC/D3 were seeded at a density of 2 × 10^5^ cells per well in 12-well plates pre-coated with 1% gelatin and allowed to adhere overnight. Subsequently, the cells were treated as indicated above (for 2, 24 and 48 h), with medium alone, vehicle control, and miR-9-5p (at concentrations of 5 nM and 10 nM). Transfection of hCMEC/D3 with the miRNA mimics was achieved using Lipofectamine RNAiMAX transfection reagent (ThermoFisher). Then, cell number and viability were assessed via a trypan blue exclusion assay. The total cell count (comprising both viable and non-viable cells) was determined utilizing an automated cell counter (Countess, ThermoFisher).

### Cell migration assay

A wound healing assay was conducted in hCMEC/D3. Briefly, cells were seeded at a density of 2 × 10^4^ cells per well in a 2-well insert with a standardized width of 500 μm (Ibidi, Gräfelfing, Germany) positioned within the wells of a 12-well culture plate pre-coated with 1% gelatin. Following an overnight incubation, the inserts were carefully removed, resulting in the creation of a uniform cell-free region within each well. Subsequently, the cells were subjected to treatment (0 to 24 h) with vehicle control and miR-9-5p (10 nM) as indicated above.

### Identification of angiogenesis-related protein expression

The expression patterns of proteins associated with angiogenesis were assessed utilizing the Proteome Profiler™ Human Angiogenesis Antibody Array Kit (R&D Systems, Minneapolis, MN, USA). Briefly, hCMEC/D3 were treated as detailed earlier with either vehicle control or miR-9-5p (10 nM) for 48 hours. After treatment, cells were lysed (in 1% Igepal CA-630, 20 mM Tris-HCl, pH 8.0, 137 mM NaCl, 10% glycerol, 2 mM EDTA, 10 μg/mL Aprotinin, 10 μg/mL Leupeptin, and 10 μg/mL Pepstatin) and subsequently used for proteome array experiment following the manufacturer’s protocol.

To quantify the targeted proteins’ concentration, the chemiluminescent signal’s intensity was captured utilizing radiographs. The obtained radiograph was then scanned utilizing a transmission-mode scanner, and the resultant image file was analyzed using ImageJ software for quantification (plugin: Protein Array Analyzer, NIH, USA).

### Protein-protein interaction (PPI) network construction and cluster analysis

Protein-protein interaction (PPI) networks of differentially expressed proteins identified from the angiogenesis-related proteomic array analysis were constructed using the Search Tool for the Retrieval of Interacting Genes (STRING) online tool (https://string-db.org) [29]. The list comprising differentially expressed angiogenic proteins was uploaded onto the platform, and the network was filtered to exhibit interactions solely with a high confidence score (≥ 0.7). To enhance the visualization of the PPI network and elucidate additional biological processes and molecular functions, a second shell comprising ten interacting proteins was incorporated based on known protein-protein interactions. After network establishment, pertinent gene ontology terms encompassing biological processes alongside Kyoto Encyclopedia of Genes and Genomes (KEGG) pathways, were delineated.

### VCAM-1 expression analysis following TNF-α **stimulation**

We wanted to determine if the effect of hsa-miR-9-5p results from global endothelial dysfunction. To achieve this, we quantified vascular cell adhesion molecule 1 (VCAM-1) protein levels in cell culture supernatants of hCMEC/D3 after stimulation with tumor necrosis factor-α (TNF-α) was conducted utilizing a commercially available human VCAM-1/CD106 DuoSet ELISA kit (R&D Systems). Briefly, hCMEC/D3 were treated with recombinant human TNF-α (R&D Systems) at a final concentration of 10 ng/mL for 16 hours. The control groups received an equivalent volume of fresh supplemented EGM-2 growth medium. Subsequently, the amount of bound VCAM-1 was assessed employing a colorimetric horseradish peroxidase (HRP) enzymatic reaction, with absorbance measurements performed at wavelengths of 450 nm and 540 nm utilizing a FLOUstar Omega microplate reader (BMG Labtech, Offenburg, Germany)

## Statistical analysis

Cell proliferation and viability assays were conducted in triplicate. Statistical comparisons between control (scramble miRNA) and hsa-miR-9-5p (5 nM and 10 nM) treatment groups across 2, 24, and 48 hours were performed using two-way ANOVA, followed by Tukey’s post hoc test. In the wound healing assay, the percentage of wound closure at 0-, 16-, and 24-hours was analyzed using repeated measures ANOVA with Bonferroni correction. The intensity of chemiluminescent signals from the Proteome Profiler™ used normalized log-transformed data and differences were assessed using the unpaired Student’s t-test. Proteins with log_2_fold (L_2_FC) > 0.5 were further analyzed. Values are presented as mean ± SD. *p*<0.05 was set as the level of significance. Graphs and statistical analyses were conducted using GraphPad Prism (GraphPad Software V10.2.2, Boston, MA, USA).

## Results

The viability of hCMEC/D3 remained unaltered following treatment with miR-9-5p at both 5 nM and 10 nM concentrations (Figure 1A). In contrast, the proliferation of hCMEC/D3 cells treated with miR-9-5p exhibited a significant dose-dependent reduction compared to controls at 24 and 48 hours (Figure 1B) (*p* < 0.05).

**Figure 1.**
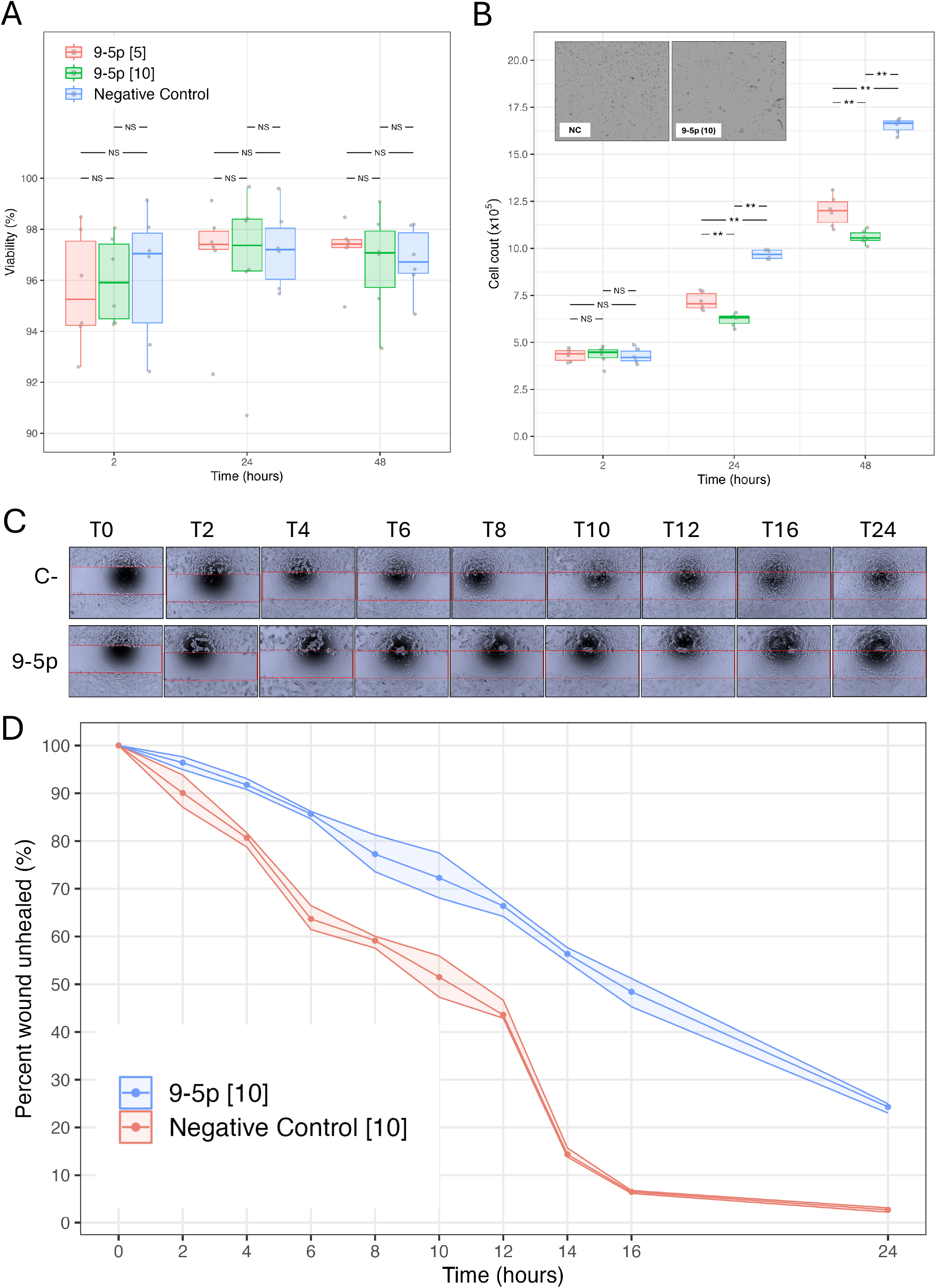
hsa-miR-9-5p reduces key angiogenic parameters in hCMEC/D3. **A)** Cell viability analysis of human cerebral microvascular endothelial cells (hCMEC/D3) exposed (2, 24 and 48 h) to mirVana™ miRNA Mimic, Negative Control (NC) (light blue), or two concentrations (5 nM, pink; and 10 nM, green) of hsa-miR-9-5p. **B)** Cell proliferation in cells treated as in A. Insert include representative images of cells exposed to NC or hsa-miR-9-5p (10 nM) after 48 h of incubation. **C)** Representative images of cell migration using wound healing assay in cells treated (T0 to T24 h) with negative control or hsa-miR-9-5p (10 nM, 9-5p). Red line indicates location of the insert that was removed at the beginning of the experiment in 100% confluent cells. Magnification 10X. **D)** Estimation of wound unhealed representing cell migration. Experimental variability is presented as shaded area in the respective group. NC: negative control, NS, non-significant differences. \*\**p*<0.001.

To delve further into the influence of hsa-miR-9-5p on cell migration, we chose the 10 nM dose for analysis. We found that hsa-miR-9-5p treatment resulted in a substantial decrease in cell migration compared to the control group (Figure 1C) (*p* < 0.05). Notably, approximately only 50% and 70% gap closure were observed by 16- and 24-hours post-treatment, significantly lower than the control group that achieved 90% and 95%, respectively (Figure 1C and 1D). These results suggest that hsa-miR-9-5p inhibits the pro-angiogenic capacity of human cerebral microvascular endothelial cells.

To support this hypothesis, we conducted an angiogenesis proteomic array, which identified 47 angiogenesis-related proteins differentially expressed in hCMEC/D3 cell lysates treated with miR-9-5p (10 nM) compared to controls (Figure 2A and 2B). Among these, 19 proteins with a log_2_fold change greater than 0.5 were selected for further analysis. PPI networks were then identified using the STRING tool (Figure 2C), with a high confidence setting (0.70). The alterations in cell proliferation/migration were found to be associated with changes in protein expression in hCMEC/D3.

**Figure 2.**
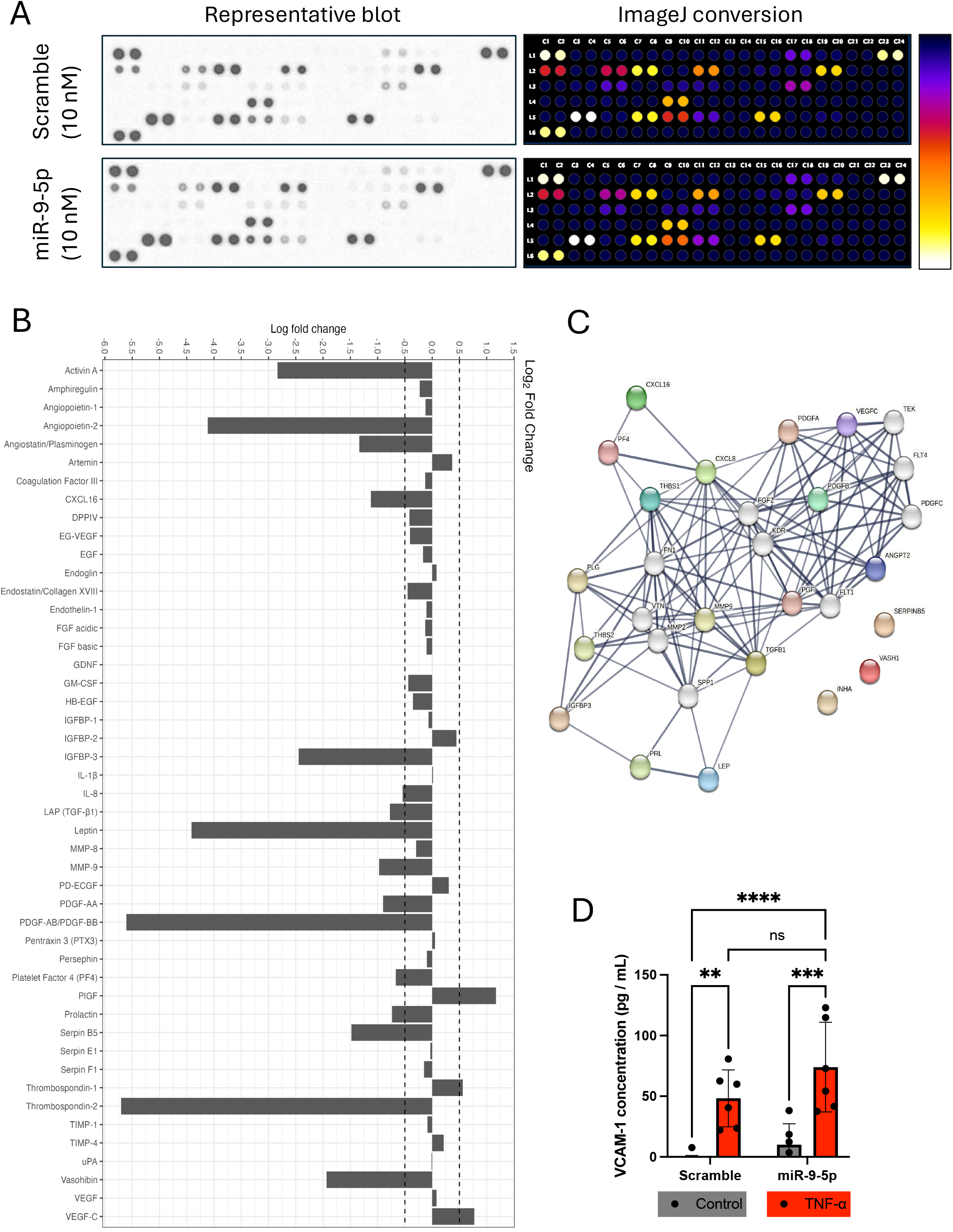
hsa-miR-9-5p modulates expression of angiogenic regulators without affecting TNF-α mediated activation of hCMEC/D3. **A)** Representative image of the Proteome Profiler™ Human Angiogenesis Antibody Array in hCMEC/D3 treated (48 h) with mirVana™ miRNA Mimic, Negative Control (Scramble), or hsa-miR-9-5p (10 nM). Left image densitometry of the representative blots. Right images represent ImageJ conversion in arbitrary color scale (white to blue, representing low and high expression respectively). **B)** Log2 fold of change of differentially expressed proteins. Scatter line represent 0.5 Log fold of change. **C)** Protein-protein interaction (PPI) network construction and cluster analysis as described in methods section. **D)** VCAM-1 protein levels in cell culture supernatants of hCMEC/D3 after stimulation with tumor necrosis factor-α (TNF-α, of 10 ng/mL for 16 hours) in presence of hsa-miR-9-5p (10 nM). The control groups received an equivalent volume of fresh supplemented EGM-2 growth medium with and without scramble construct. VCAM-1: vascular cell adhesion molecule 1, NS, non-significant differences. \*\**p*<0.01. \*\*\**p*<0.001; \*\*\*\**p*<0.0001.

The differentially expressed positive regulators of endothelial cell migration were platelet derived growth factor B (PDGFB, L_2_FC -5.6), vascular endothelial growth factor C (VEGFC, L_2_FC 0.77) and plasminogen (PLG, L_2_FC -1.34). Vasohibin 1 (VASH1, L_2_FC -1.94) and angiopoietin 2 (ANGPT2, L_2_FC -4.11) were identified as negative regulators of endothelial cell migration. Regarding endothelial cell proliferation, VEGFC, placental growth factor (PLGF or PGF – as denoted in the PPI network, Fig. 2C, L_2_FC 1.17) and PDGFB were identified as positive regulators, whereas prolactin (PRL, L_2_FC -0.74), thrombospondin 1 (THBS1, L_2_FC 0.56) and VASH1 were identified as negative regulators. The PPI networks and KEGG pathways previously associated with preeclampsia, including the hypoxia inducible factor 1 (HIF-1) signalling pathway and focal adhesions, were also identified. All differentially expressed proteins with their respective L_2_FC changes, along with the list of identified biological processes and KEGG pathways can be found in the supplementary materials (Table S1-S3).

Finally, we sought to determine whether these changes represent endothelial dysfunction or solely affect angiogenesis processes by investigating the response of hCMEC/D3 cells treated with vehicle control and miR-9-5p (10nM) to TNF-α stimulation. hCMEC/D3 treated with scramble and hsa-miR-9-5p (10nM) without TNF-α stimulation were used as controls. Expression of VCAM-1, a marker of endothelial cell activation, was analyzed in the culture supernatants. Results indicate that treatment with miR-9-5p did not impair the capacity of hCMEC/D3 to respond to pro-inflammatory TNF-α (Figure 2D).

## Discussion

We found that hsa-miR-9-5p downregulates angiogenesis-related processes, including endothelial cell proliferation and migration, without affecting TNFα-mediated endothelial activation. This finding is supported by the altered expression levels of angiogenesis-related proteins in hCMEC/D3 treated with miR-9-5p and preserved responses to VCAM-1 induction and expression. These findings suggest that the cargo within elevated circulating syncytiotrophoblast-derived extracellular vesicles may contribute to the brain endothelial dysfunction observed in preeclampsia.

Severe preeclampsia is associated with a range of maternal brain complications, including posterior reversible encephalopathy syndrome, white matter abnormalities, cerebrovascular disorders, and stroke, highlighting its significant impact on neurological health [30], and constituting one of the leading cause of maternal mortality associated to preeclampsia [1-3]. Underlying mechanisms of cerebrovascular complications in preeclampsia involve vasogenic oedema, cerebral vasoconstriction, and disruption of the blood-brain barrier (BBB), which potentially leads to eclampsia (i.e., seizures) [31]. How these defects are generated is still under investigation, with increasing number of evidence showing the participation of circulating factors present in the maternal circulation, which are likely derived from the placenta [32]. Among them, some reports [33-35] indicate that placental derived extracellular vesicles, such as STB-EVs, may contribute to cerebrovascular manifestations. hsa-miR-9-5p is one of the m/lSTB-EV cargo significantly overexpressed in preeclampsia [26], suggesting a potentially significant role in triggering cerebral endothelial dysfunction and influencing recovery due to its effects as an angiogenic modulator [27, 28]. We speculate that by targeting angiogenesis, miR-9-5p may contribute to the cerebrovascular alterations observed in preeclampsia, impairing cerebral autoregulation [36-38], and potentially affecting cerebral blood flow. Further studies are required to evaluate the *in vivo* role of miR-9-5p in those cerebrovascular parameters.

Furthermore, EVs isolated from the plasma of women with preeclampsia or from hypoxic placentae have been shown to disrupt the BBB both *in vitro* [33, 34] and *in vivo* [33, 35]. Notably, the *in vitro* reported models used the same human cerebral microvascular endothelial cells (i.e., hCMEC/D3) described in our study. Our findings add to this understanding by demonstrating that cargo within m/lSTB-EVs, such as hsa-miR-9-5p, may impair the angiogenic capacity of these cells.

The mechanisms by which hsa-miR-9-5p reduces cell proliferation and migration and modulates angiogenic protein regulators in cerebral microvascular endothelial cells, remain unclear. Indirect evidence from choroidal melanoma suggests that hsa-miR-9-5p inhibits proliferation, migration, and invasion by suppressing the transcription and translation of BRAF mRNA [39]. Additionally, in chronic cerebral hypoperfusion (CCH) rats, inhibiting hsa-miR-9-5p with antagomirs rescues learning and memory impairments, synaptic plasticity, dendritic spines, cholinergic neurons, oxidative stress levels, and neuronal loss [40]. Future studies should focus on elucidating the specific downstream targets and signaling pathways of hsa-miR-9-5p in hCMEC/D3 cells to better understand its role in regulating brain angiogenesis.

Contrary to our findings, some previous studies suggest that hsa-miR-9-5p has a pro-angiogenic role in human umbilical vein endothelial cells (HUVECs) by upregulating the VEGF/VEGFR2 signaling pathway or targeting the CXC chemokine receptor-4 (CXCR4) [27, 28]. This underscores the importance of considering cell type specificity when investigating microRNA function. In our study, we used hCMEC/D3, a human microvascular endothelial cell line and a well-established model for studying the BBB with a maternal origin [20], whereas the referenced studies used fetal endothelial cells from the macro-circulation. It’s important to note that the endothelium of the micro and macro circulation are physiologically distinct [41].

One of the primary regulators of angiogenesis is the VEGF and VEGFR protein families [42]. Our results show that hCMEC/D3 cells treated with hsa-miR-9-5p overexpressed PLGF and VEGFC, two pro-angiogenic proteins within the VEGF family [42]. However, PDGFB, a positive regulator of endothelial cell proliferation and migration, was significantly downregulated in these cells (L_2_FC -5.60). The PPI network also identified VEGFR2 as a key interacting protein, playing a critical role in endothelial cell proliferation and migration. Notably, the activation of VEGFR2 (also known as kinase insert domain receptor, KDR) is crucial for angiogenesis [43]. This suggests that the functional modulation of the VEGF/VEGFR signaling pathway may contribute to the anti-angiogenic effects of hsa-miR-9-5p observed in our study. However, the dynamic interactions of these proteins require further investigation.

In addition to its pro-angiogenic role, VEGFR2 in brain endothelial cells may contribute to the functional stabilization of the BBB [18, 20, 44]. We propose that the overexpression of VEGFC and PLGF in hCMEC/D3 cells treated with hsa-miR-9-5p may also modulate VEGFR2 activation, a key feature observed in in vitro studies of the BBB in preeclampsia [20]. Furthermore, the preserved response of hCMEC/D3 cells to TNF-α stimulation in the presence of hsa-miR-9-5p suggests that the observed effects specifically impair angiogenesis rather than cause overall endothelial dysfunction.

Our work raises several new questions. For example, at what gestational age do circulating levels of hsa-miR-9-5p begin to rise? In our previous report, we identified elevated levels of hsa-miR-9-5p in STB-EVs from EOPE pregnancies (i.e., < 34 weeks of gestation) [26], suggesting that this increase may correspond with early placental alterations observed in this condition [45]. This raises the intriguing possibility that hsa-miR-9-5p could serve as a biomarker linking placental and cerebrovascular changes. We encourage future cohort studies to explore this potential.

Another question is whether hsa-miR-9-5p also affects endothelial cells in other critical organs involved in maternal pregnancy adaptations, such as the kidneys. The PPI network identified the regulation of glomerular vasculature as a key biological process. Previous studies have reported glomerular endothelial cell swelling (glomerular endotheliosis) as a feature of preeclampsia [46]. Therefore, investigating the potential role of hsa-miR-9-5p, carried within m/lSTB-EVs, in contributing to endothelial dysfunction in renal glomeruli is warranted.

This study’s strengths include the use of a human cerebral endothelial cell line derived from a woman with a history of epilepsy [47], which may provide a relevant model for brain endothelial dysfunction in preeclampsia. Additionally, we employed advanced techniques such as proteomic array analysis and PPI networks to deepen our understanding of the molecular mechanisms underlying these effects. However, we recognize that *in vitro* models cannot fully replicate the complexity of the in vivo microenvironment, which may limit the translation of our findings to *in vivo* settings.

In conclusion, our study highlights the significant role of hsa-miR-9-5p, which is highly expressed in EOPE m/lSTB-EVs, in affecting key parameters related to brain angiogenesis. This novel finding offers valuable insights into how STB-EV cargo impairs cerebrovascular function in preeclampsia. Our results pave the way for future research to determine whether hsa-miR-9-5p could serve as a potential biomarker for the early detection of cerebrovascular complications in preeclampsia. Further understanding of hsa-miR-9-5p’s role in STB-EVs could advance broader research into EV-mediated disruption of angiogenesis and vascular integrity.

## Conflict of interest

None to declare.

## Author contribution

MV conceptualized the study and lead the research team. PL, TA, MR and NA performed the experiments in this manuscript. GDJ performed statistical analysis and data visualization. PL, TA and CE wrote the draft of the manuscript. MV and ASC supervised the project. All co-authors approved the final version of this manuscript.

## Acknowledgement

NA acknowledges support by research grants from the British Heart Foundation (BHF) Centre of Research Excellence, Oxford (RE/13/1/30181 and RE/18/3/34214) and a British Heart Foundation Intermediate Fellowship (FS/IBSRF/22/25110). The Tripartite Immunometabolism Consortium, Novo Nordisk Foundation (NNF15CC0018486 and NNF20SA0064144). The views expressed are those of the author(s) and not necessarily those of the National Health Service, the National Institutes of Health Research, or the Department of Health. CE is funded by Fondecyt 1240295.

## Abbreviations

(ANGPT2): Angiopoietin 2
(BBB): Blood-brain barrier
(EOPE, < 34 weeks): Early-onset preeclampsia
(EVs): Extracellular vesicles
(KDR): Kinase insert domain receptor
(> 200 nm, m/lSTB-EVs): Medium/large STB-EVs
(miRNAs): MicroRNAs
(PLGF or PGF): Placental growth factor
(PE): Preeclampsia
(PRL): Prolactin
(< 200 nm, sSTB-EVs): Small STB-EVs
(STB-EVs): Syncytiotrophoblast
(STB-EVs): Syncytiotrophoblast-derived extracellular vesicles
(THBS1): Thrombospondin 1
(TNF-α): Tumor necrosis factor-α
(BRAF): V-raf murine sarcoma viral homolog B1
(VCAM-1): Vascular cell adhesion protein 1
(VEGFC): Vascular endothelial growth factor type C
(VASH1): Vasohibin 1
(VEGFR2): VEGF receptor 2

## Supplementary tables

S1 - Log fold changes of differentially expressed angiogenesis-related proteins in hCMEC/D3 treated with miR-9-5p (10 nM)

S2 - Gene ontology terms (biological processes) associated with established PPI network

S3 - KEGG pathways associated with established PPI network

## Notes

### Competing Interest Statement

The authors have declared no competing interest.

