## supplementary materials for "hsa-miR-9-5p highly expressed in syncytiotrophoblast-derived extracellular vesicles from early-onset preeclampsia impairs cerebral microvascular endothelial cell pro-angiogenic capacity"

| **Under-expressed proteins** | **LFC** | **Over-expressed proteins** | **LFC** |
| --- | --- | --- | --- |
| Thrombospondin-2 (THBS2) | -5.70 | Placental growth factor (PlGF/PGF) | 1.17 |
| Platelet-derived growth factor-BB (PDGFB) | -5.60 | Vascular endothelial growth factor-C (VEGFC) | 0.77 |
| Leptin (LEP) | -4.41 | Thrombospondin-1 (THBS1) | 0.56 |
| Angiopoeitin-2 (ANGPT2) | -4.11 | Insulin-like growth factor binding protein-2 (IGFBP2) | 0.44 |
| Activin-A (INHBA) | -2.84 | Artemin (ARTN) | 0.37 |
| Insulin-like growth factor binding protein-3 (IGFBP3) | -2.45 | Platelet-derived endothelial cell growth factor (PD-ECGF) | 0.30 |
| Vasohibin (VASH1) | -1.94 | TIMP metallopeptidase inhibitor 4 (TIMP4) | 0.21 |
| Serpin B5 (SERPINB5) | -1.48 | Endoglin (ENG) | 0.08 |
| Plasminogen (PLG) | -1.34 | Vascular endothelial growth factor (VEGF) | 0.08 |
| Chemokine (C-X-C motif) ligand 16 (CXCL16) | -1.12 | Pentraxin 3 (PTX3) | 0.05 |
| Matrix metalloproteinase-9 (MMP9) | -0.97 | Interleukin 1β (IL-1β) | 0.02 |
| Platelet-derived growth factor-AA (PDGFA) | -0.90 |  |  |
| Transforming growth factor beta-1 (TGFB1) | -0.78 |  |  |
| Prolactin (PRL) | -0.74 |  |  |
| Platelet factor 4 (PF4) | -0.67 |  |  |
| Interleukin 8 (CXCL8) | -0.54 |  |  |
| Endostatin (COL18A1) | -0.45 |  |  |
| Granulocyte-macrophage colony-stimulating factor (GM-CSF) | -0.44 |  |  |
| Dipeptidyl peptidase-4 (DPPIV) | -0.42 |  |  |
| Endocrine gland derived vascular endothelial growth factor  (EG-VEGF) | -0.41 |  |  |
| Heparin-binding epidermal growth factor-like growth factor (HBEGF) | -0.35 |  |  |
| Matrix metalloproteinase-8 (MMP8) | -0.29 |  |  |
| Amphiregulin (AREG) | -0.23 |  |  |
| Epidermal growth factor (EGF) | -0.17 |  |  |
| Serpin F1 (SERPINF1) | -0.15 |  |  |
| Coagulation factor III (F3) | -0.13 |  |  |
| Fibroblast growth factor acidic (FGF1) | -0.13 |  |  |
| Angiopoeitin-1 (ANGPT1) | -0.12 |  |  |
| Endothelin-1 (EDN1) | -0.10 |  |  |
| Fibroblast growth factor basic (FGF2) | -0.10 |  |  |
| Persephin (PSPN) | -0.09 |  |  |
| TIMP metallopeptidase inhibitor 1 (TIMP1) | -0.09 |  |  |
| Insulin-like growth factor binding protein-1 (IGFBP1) | -0.07 |  |  |
| Serpin E1 (SERPINE1) | -0.03 |  |  |
| Plasminogen activator, urokinase receptor (PLAUR) | -0.01 |  |  |
| Glial cell line derived neurotrophic factor (GDNF) | -0.001 |  |  |

S1: Log fold changes of differentially expressed angiogenesis-related proteins in hCMEC/D3 treated with miR-9-5p (10 nM)

| **Biological processes** | **Strength** |
| --- | --- |
| Negative regulation of phosphatidylinositol biosynthetic process | 2.83 |
| Positive regulation of metanephric mesenchymal cell migration by PDGF-receptor-beta signalling pathway | 2.66 |
| Tie signalling pathway | 2.53 |
| Positive regulation of mast cell chemotaxis | 2.43 |
| Regulation of lymphangiogenesis | 2.43 |
| Positive regulation of follicle-stimulating hormone secretion | 2.35 |
| Paracrine signalling | 2.29 |
| Regulation of blood vessel remodelling | 2.29 |
| Negative regulation of vasoconstriction | 2.23 |
| Regulation of hyaluronan biosynthetic process | 2.23 |
| VEGF signalling pathway | 2.21 |
| Negative regulation of fibrinolysis | 2.20 |
| Induction of positive chemotaxis | 2.16 |
| Ovulation from ovarian follicle | 2.13 |
| Hyaluronan catabolic process | 2.13 |
| Regulation of positive chemotaxis | 2.12 |
| Cellular response to UV-A | 2.09 |
| Positive regulation of positive chemotaxis | 2.04 |
| Positive regulation of protein autophosphorylation | 2.02 |
| VEGF-receptor signalling pathway | 2.01 |
| Regulation of endothelial cell chemotaxis | 2.01 |
| Negative regulation of blood vessel endothelial cell migration | 1.99 |
| Positive regulation of phosphatidylinositol 3-kinase activity | 1.99 |
| Glomerulus vasculature development | 1.99 |
| Positive regulation of endothelial cell chemotaxis | 1.99 |
| Surfactant homeostasis | 1.96 |
| Positive regulation of blood vessel endothelial cell migration | 1.95 |
| Regulation of macrophage differentiation | 1.95 |
| Positive regulation of fibroblast migration | 1.93 |
| Regulation of blood vessel endothelial cell migration | 1.93 |

**S2: Gene ontology terms (biological processes) associated with established PPI network**

| **KEGG pathways** | **Strength** |
| --- | --- |
| Bladder cancer | 1.83 |
| Malaria | 1.77 |
| Focal adhesion | 1.66 |
| EGFR tyrosine kinase inhibitor resistance | 1.64 |
| Rap1 signalling pathway | 1.61 |
| ECM-receptor interaction | 1.59 |
| Melanoma | 1.58 |
| AGE-RAGE signalling pathway in diabetic complications | 1.55 |
| PI3K-Akt signalling pathway | 1.52 |
| Ras signalling pathway | 1.52 |
| Rheumatoid arthritis | 1.52 |
| MAPK signalling pathway | 1.45 |
| Proteoglycans in cancer | 1.45 |
| Prostate cancer | 1.45 |
| p53 signalling pathway | 1.45 |
| Fluid sheer stress and atherosclerosis | 1.42 |
| Gap junction | 1.37 |
| MicroRNAs in cancer | 1.33 |
| Relaxin signalling pathway | 1.33 |
| Choline metabolism in cancer | 1.33 |
| Renal cell carcinoma | 1.32 |
| Transcriptional misregulation in cancer | 1.30 |
| HIF-1 signalling pathway | 1.30 |
| Amoebiasis | 1.30 |
| Glioma | 1.28 |
| Phospholipase D signalling pathway | 1.27 |
| JAK-STAT signalling pathway | 1.24 |
| Cytokine-cytokine receptor interaction | 1.23 |
| Regulation of actin cytoskeleton | 1.21 |
| Pathways in cancer | 1.16 |

S3: KEGG pathways associated with established PPI network
